# *Bifidobacterium pseudocatenulatum* capsular exopolysaccharide enhances systemic anti-tumour immunity in pre-clinical breast cancer

**DOI:** 10.1101/2024.09.23.614466

**Authors:** Christopher A. Price, Alicia Nicklin, Magdalena Kujawska, Todor T. Koev, Nilda Ilker, Wesley J. Fowler, Alastair M. McKee, Luke Mitchell, Mitchel Rowe, James A.G.E. Taylor, Christopher J. Benwell, Sally A. Dreger, Julia Mueller, Lindsay J. Hall, Stephen D. Robinson

## Abstract

Gut microbes have merged as powerful regulators of cancer responses, with *Bifidobacterium* species and strains playing a key role in promoting anti-tumour immunity. While they represent promising candidates for cancer therapeutics, the specific underlying microbial mechanisms driving their efficacy remains poorly understood. In this study, we demonstrate the broad potential of *Bifidobacterium* species to inhibit breast cancer progression across multiple pre-clinical mouse models. We identify a novel strain, *Bifidobacterium pseudocatenulatum* 210, which induces systemic anti-tumour immunity and enhances responses to standard-of-care therapies via its cell surface capsular exopolysaccharide (EPS). *B. pseudocatenulatum* 210 EPS promotes dendritic cell activation and increases systemic cDC1 infiltration, leading to robust CD8^+^ T cell-mediated anti-tumour activity. Our findings position *Bifidobacterium* EPS as a novel class of therapeutic compounds with significant potential for cancer treatment.

## Introduction

One of the key challenges of cancer therapy is inconsistent patient response to standard of care treatments. Host-intrinsic factors, particularly the gut microbiome, are now recognised as significant contributors to resistance mechanisms that mediate these variable therapeutic outcomes (*1, 2*). The gut microbiome’s role in regulating cancer progression has been demonstrated in both mice (*3-5*) and humans (*6-8*), where specific bacterial species and strains can both protect against cancer progression and enhance responses to standard of care therapy (*3, 9, 10*). However, while the concept of anti-tumour activity between intestinal bacteria is established, most studies to date focus on immunogenic tumours (e.g., melanoma, lung cancer) (*11-14*) and gastrointestinal cancers like colorectal cancer (*9, 15*). The potential influence of gut bacteria on extra-intestinal non-immunogenic indications, like breast cancer, remains poorly characterised.

Beneficial bacteria, such as *Bifidobacterium* and *Lactobacillus*, are known to inhibit solid tumour types in pre-clinical models (*9, 16, 17*). Several species of *Bifidobacterium*, including *Bifidobacterium bifidum (15)*, *Bifidobacterium breve (18, 19)*, and *Bifidobacterium pseudolongum (10)* have been linked with anti-tumour immunity. Human studies further support these findings, showing positive correlations between abundances of various *Bifidobacterium* species and improved patient outcomes across multiple cancer types (*11, 14*). Collectively, this growing body of evidence supports *Bifidobacterium*-based therapeutic interventions to treat cancer.

Despite these advances, microbiome-based cancer therapeutics have often lacked a clear mechanistic understanding of the microbial-derived active compounds responsible for their beneficial effects. While, microbial metabolites have been most described (*9, 10, 20, 21*), emerging research highlights the role of microbial structural compounds, such as peptidoglycan and exopolysaccharides (EPS), in modulating protective responses (*15, 22*). These exopolysaccharides, composed of polymeric sugar chains with diverse glycosyl content and structure, can drive specific host immune pathways (*23, 24*). A seminal study by *Vétizou et al*., (*3*) demonstrated *Bacteroides fragilis* polysaccharide A (PSA) mediates anti-tumour immunity response when combined with immune checkpoint inhibitors. More recently, *Sharma et al.*, (*22*) showed that a strain of *Lactobacillus plantarum* surface polysaccharide promotes anti-tumour immunity through enhanced activity of macrophage iron sequestration pathways, skewing tumour-associated macrophages to an anti-tumour CD8^+^-permissive state. However, the role of *Bifidobacterium-*derived EPS in stimulating cancer immunity *in vivo* has yet to be explored.

In this study, we demonstrate that several species of *Bifidobacterium* (*Bifidobacterium bifidum*, *Bifidobacterium reuteri*, and *Bifidobacterium pseudocatenulatum*) exhibit therapeutic efficacy in pre-clinical breast cancer models. We show that administration of *B. pseudocatenulatum* 210 reduces primary tumour burden across models of the major breast cancer subtypes and enhances responses to standard-of-care chemotherapy and immunotherapy. We demonstrate a novel mechanism whereby *B. pseudocatenulatum* 210 capsular EPS mediates drives systemic anti-tumour immunity from the gut. The EPS induces CD8^+^ T cell-dependent anti-tumour immunity via dendritic cells (DCs) activation, promoting the maturation of CD80^+^CD86^+^ DCs and increasing systemic infiltration of CD8^+^ T cell specific cDC1 cells. These 210 EPS-programmed DCs enhance polarisation and activation of CD8^+^ T cells at systemic sites and, crucially, increase the fraction of IFNγ^+^TNFα^+^ poly-functional CD8^+^ T cells at the primary tumour. We demonstrate that 210-derived EPS can be isolated and used therapeutically independently of any live parental *B. pseudocatenulatum* cells, positioning *Bifidobacterium* EPS as a promising new cancer immunotherapy compound.

## Methods

### Mice

C57 BL/6 mice were purchased and maintained in-house at the Disease Modelling Unit at the University of East Anglia under project licence code PP8873233. BALB/C mice were purchased from Charles River and maintained in-house. Animals used throughout were age 8-12 weeks and were randomly mixed between cages prior to experiment onset. All animal experiments were performed in accordance with UK Home Office regulations and the European Legal Framework for the Protection of Animals used for Scientific Purposes (European Directive 86/609/EEC).

### Orthotopic breast tumour growth experiments

Syngeneic breast cancer cells were injected in 50μl of a 1:1 mixture of phosphate buffered saline (PBS) and Matrigel (Corning Life Sciences, Corning, USA) into the left inguinal mammary fat pad of age-matched female mice. PyMT-BO1, E0771 and 4T1 cells were each injected 1×10^5^ cells and BRPKp110 cells were injected at 5×10^5^ cells. The experimental duration for each breast model used is outlined in the associated figures. In situ tumour volumes were measured thrice weekly with digital callipers from the onset of a palpable tumour, using the formula: length x width^2^ x 0.52 (*25*).

### *In vivo* experimental interventions (orthotopic experiments)

Animals were orally administered thrice weekly with live *Bifidobacterium* strains (1×10^10^ CFU/200μl) or isolated EPS (80μg/200μl) from the onset of a palpable tumour to endpoint. For chemotherapy experiments, cyclophosphamide (Sigma) was administered by intraperitoneal injection at 100mg/kg on days 10 and 17. For checkpoint immunotherapy experiments, tumour-bearing animals were intraperitoneally administered 20mg/kg anti-PD-1 mAb (Clone J43, BioXCell, New Hampshire, USA) or matched isotype control on days 7, 10, 13. Cellular depletions were induced through intraperitoneal injection of 400μg anti-CD8-α (clone 2.43, BioXCell) or matched isotype control mAb one day prior to *Bifidobacterium* administration (day 9), followed by 200μg injections thereafter on days 13, 16, and 19. Depletion was verified by flow cytometry of primary tumour immune cells. For microbiome depletion, BRPKp110 tumour-bearing animals were treated with antibiotics by oral gavage (200μl in water vehicle) on day 3, 6 and 8 prior to therapeutic intervention. The antibiotic cocktail contained 1mg/ml Amphotericin B, 25mg/ml Vancomycin, 50mg/ml Neomycin and 50mg/ml Metronidazole (all purchased from Sigma). Animal drinking water was also supplemented with 1mg/ml Ampicillin (Sigma). For adoptive cell transfer experiments, 1×10^6^ bone marrow-derived dendritic cells (BMDCs) were cultured with either 5×10^6^ CFU live 210 cells, 80μg of EPS derived from *B. pseudocatenulatum* 210 or *B. longum* B71, or vehicle control (PBS). Treated BMDCs were adoptively transferred by intravenous injection to naïve BRPKp110 tumour-bearing animals on days 10, 14, and 18.

### PyMT spontaneous tumour growth experiments

MMTV-PyMT (B6.FVB-Tg(MMTV-PyVT)634Mul/LellJ) mice were obtained from Jackson Laboratory (stock 022974) on a congenic C57BL/6 background. Female mice heterozygous for the PyMT transgene were used for tumour growth studies. Mice were palpated for tumours and treated twice weekly with oral administrations of *B. pseudocatenulatum* 210 from 8 weeks of age, with the first palpable tumours being observed from 12 weeks. Tumour growth was measured using digital calipers and animals were sacrificed once tumours reached 1cm^3^ or at the 24-week experimental endpoint.

### Germ free colonisation experiments

Germ free mice were maintained in isolators and orally administered 1×10^10^ CFU of *B. pseudocatenulatum* 210 or vehicle control (PBS). Animals were sacrificed at 6, 24, and 48 hours post administration and gut contents were isolated from the upper and lower colon, caecum, and small intestine in sterile MSC class II cabinets. Gut contents were isolated at 50mg per sample, per animal, and mechanically homogenised in 1ml of PBS. 100µL of the slurry solution was then cultured on MRS-cysteine agar plates and incubated anaerobically (37°C) for 48 hours. Visible colonies were counted, and bacterial load (CFU/g) was calculated for each condition with the following: CFU/g = (no. of colonies x dilution factor) / (volume cultured x weight of faecal sample). Gut contents from PBS treated germ free animals were confirmed to be sterile at each timepoint by the same method.

### Bacterial culture

All *Bifidobacterium* strains (see supplementary Table 1) used for this study were isolated previously by the laboratory of Prof. Lindsay Hall. The strains were cultured at 37°C in MRS broth with L-cysteine (50mg/L) (Sigma) in an anaerobic chamber (Don Whitley Scientific, Bingley, UK). Strains were cultured for one week and then preserved by lyophilisation in the exponential phase of growth. Lyophilised bacteria were equally distributed across individual dosage vials which were stored at -80°C. To ensure accurate dosing, at least three vials from each lyophilisation batch were enumerated by counting CFUs on MRS agar plates across serial PBS dilutions. The mean CFU was calculated for each batch and vials were resuspended in PBS (to 1×10^10^ CFU/200μl/animal) immediately prior to experimental administrations.

For acid-killed experiments, peracetic acid pre-treatment of bacteria was performed as previously described (*26*). Briefly, lyophilised bacteria were reconstituted in 10ml of sterile PBS at a concentration of ∼1×10^10^ CFU/ml. Peracetic acid (Sigma) was added to a final concentration of 0.4% and bacteria were incubated at room temperature (RT) for 1 hour. The bacteria were washed three times in sterile PBS and then resuspended to the appropriate final concentration for animal administrations. Bacterial killing was confirmed by culturing 100μl of acid-killed bacteria on MRS agar under anaerobic conditions and validating the absence of bacterial growth compared with a non-acid treated positive control.

### Cell culture

All tumour cell lines were cultured in high glucose DMEM (Thermofisher) supplemented with 10% foetal bovine serum (FBS) (Hyclone, Thermofisher) and 100 units/ml penicillin/streptomycin (Thermofisher). Cells were seeded onto flasks coated with 0.1% porcine gelatin (Sigma) and incubated at 37°C and 5% C0_2_.

BMDCs were generated from flushed bone marrow from tibias and femurs of C57 BL/6 mice. Isolated bone marrow was treated with red blood cell lysis buffer for 5 minutes, washed twice in PBS, and cultured overnight in RPMI1640 medium (Thermofisher) supplemented with 10% FBS, 1% penicillin/streptomycin (Gibco), 20ng/ml IL-4 (R&D Systems) and 10ng/ml GM-CSF (R&D Systems). On the second day, supernatant with non-adherent cells was removed and culture media replaced. Cell culture medium was replaced again on the fifth day, and semi-adherent BMDCs were collected on the seventh day for experimental use.

### Lung histology

Harvested organs were incubated overnight in 4% paraformaldehyde (PFA) at 4°C and processed with the Leica Tissue Processor ASP-300-S (Leica Biosystems, Milton Kenes, UK). The tissues were incubated in formalin, dehydrated through increasing concentrations of ethanol (from 70% to 100%), washed in three changes of xylene (Sigma) and then embedded in paraffin (Sigma). Paraffin blocks were sectioned at 6μm using a rotary microtome (Leica Biosystems, RM2235), mounted onto positively charged glass slides (Thermofisher), and incubated overnight at 37°C. Prior to histological staining, FFPE tissue sections were deparaffinised in xylene (Sigma) and rehydrated through sequentially decreasing concentrations of ethanol (100%-70%), then into water. H&E staining was performed using a Leica ST5020 tissue multi-stainer (Leica Biosystems, Nussloch, Germany) and sections were then mounted with coverslips with Neo-Mount™ (Sigma). Images were captured using the Olympus BX60 microscope (Olympus, Southend-on-Sea, UK) with a microscope camera Jenoptik C10 and ProgRes CapturePro software v.2.10.

### Flow cytometry

Tumours and lungs were excised and mechanically homogenised using scalpels. Tumours were digested in 0.2% collagenase IV (Thermofisher) and lungs in 0.2% collagenase I (Thermofisher), with 0.01% hyaluronidase (Sigma) and 0.01% DNase I (in HBBS) at 37°C under agitation for 60 minutes (tumours) or 30 minutes (lungs). Tissues were then passed through a 70µm filter (Thermofisher) and washed in PBS before further staining. Spleens and lymph nodes were mechanically dissociated through 70µm strainers. Lymph nodes were immediately processed for staining and tumours, lungs, and spleens were resuspended in red blood cell lysis buffer (Thermofisher) for 5 minutes. Blood was collected by cardiac puncture in EDTA and washed twice in red blood cell lysis buffer prior to cell staining. Cells from all tissues were washed in PBS and 1×10^6^ cells per sample were then resuspended in FACS buffer (2% FBS in PBS) prior to further staining with extracellular antibodies.

For intracellular cytokine analysis, cells were resuspended in 200µl RPMI supplemented with 10% FBS, 50µM 2-Mercaptoethanol, 50ng/ml Phorbol 12-Myristate 13-Acetate (PMA), 750ng/ml Ionomycin and 10µg/ml Brefeldin-A (all purchased from Sigma) and incubated in a 96-well U-bottom plate (Sigma) at 37°C, 5% CO_2_ for 4 hours. Cells were blocked with Fc-receptor blocking reagent (Thermofisher) and incubated in relevant extracellular antibody (Table 1) and fixable Live/Dead Red (Invitrogen) solutions for 30 minutes (at 4°C in the dark). Cells were washed twice in FACS buffer, fixed with 4% PFA for 30 minutes, and then resuspended in FACS buffer prior to analysis. Where intracellular staining was required, cells were treated using the eBioscience™ Foxp3 / Transcription Factor Staining Buffer Set (Thermofisher), as per the manufacturer’s instructions, then stained with intracellular antibodies.

**Table 1.** List of conjugated flow cytometry antibodies.

| Target | Conjugate | Clone | Source | Category number | Dilution |
| --- | --- | --- | --- | --- | --- |
| CD45 | PerCP-Cy5.5 | 30-F11 | Invitrogen | 45-0451-02 | 1:200 |
| CD3 | APC | 17A2 | BioLegend | 100236 | 1:200 |
| CD8a | APC-Cy7 | 53-6.7 | Invitrogen | A15386 | 1:200 |
| CD44 | eFluor450 | 1M7 | Invitrogen | 48-0441-82 | 1:200 |
| CD62L | BV605 | MEL-14 | BioLegend | 104438 | 1:200 |
| NK1.1 | PE-Cy7 | PK136 | BioLegend | 108714 | 1:200 |
| IFNγ | BV711 | XMG1.2 | BioLegend | 505836 | 1:100 |
| IL-2 | PerCP-Cy5.5 | JES6-5H4 | Invitrogen | 45-7021-80 | 1:100 |
| FOXP3 | eFluor450 | FJK-16s | Invitrogen | 48-5773-82 | 1:100 |
| CD45 | BUV395 | 30-F11 | BD | 564279 | 1:200 |
| CD4 | PE | GK1.5. | Invitrogen | 12-0041-82 | 1:200 |
| FOXP3 | AF700 | FJK-16s | Invitrogen | 56-5773-82 | 1:100 |
| Granzyme B | PE-Cy7 | NGZB | Invitrogen | 25-8898-82 | 1:200 |
| CD11b | BV605 | M1/70 | BD | 563015 | 1:200 |
| Ly6C | eFluor450 | HK1.4 | Invitrogen | 48-5932-82 | 1:200 |
| Ly6G | APC-Cy7 | 1A8 | BD | 560600 | 1:200 |
| F4/80 | APC | BM8 | Invitrogen | 17-4801-82 | 1:200 |
| MHCII (I-A/I-E) | PE-Cy7 | M5/114.15.2 | Invitrogen | 25-5321-82 | 1:200 |
| CD206 | FITC | CO68C2 | BioLegend | 141704 | 1:200 |
| CD11c | BUV395 | HL3 | BD | 564080 | 1:200 |
| CD103 | BV711 | 2E7 | BioLegend | 121435 | 1:200 |
| PD-1 | PE-Cy7 | 29F.1A12 | BioLegend | 135216 | 1:200 |
| CD80 | Super Bright 600 | GL1 | Invitrogen | 63-0862-82 | 1:200 |
| CD86 | BV421 | 16-10A1 | BD | 562611 | 1:200 |
| CD107a | eFluor450 | 104B | Invitrogen | 48-1071-82 | 1:100 |
| GFP | PE-Cy7 | FM264G | BioLegend | 338014 | 1:100 |
| IL-4 | PerCP-Cy5.5 | 11B11 | BioLegend | 504123 | 1:100 |
| TNFα | PE | MP6-XT22 | BioLegend | 506306 | 1:100 |

Data collection was conducted on the BD LSR Fortessa cell analyser and analysed using FlowJo software (BD). All samples were initially gated using FSC-A vs. FSC-H to identify single cells (Singlets), which were then gated for Live/Dead Red negative. Major immune populations were identified using the marker profiles outlined in Table 2.

**Table 2.** List of flow cytometry gating strategies for the identification of immune cell populations.

| Population | Gating strategy |
| --- | --- |
| Lymphoid cells | Singlet, Live, CD45+, CD3+ |
| T helper cells | Singlet, Live, CD45+, CD3+, CD4+, CD8- |
| **CD8<sup>+</sup> T cells** | Singlet, Live, CD45+, CD3+, CD4-, CD8+ |
| **CD8<sup>+</sup> effector memory** | Singlet, Live, CD45+, CD3+, CD4-, CD8+, CD62L-, CD44+ |
| **CD8<sup>+</sup> central memory** | Singlet, Live, CD45+, CD3+, CD4-, CD8+, CD62L+, CD44+ |
| **Naïve CD8** | Singlet, Live, CD45+, CD3+, CD4-, CD8+, CD62L+, CD44- |
| **Treg cells** | Singlet, Live, CD45+, CD3+, CD4+, CD8-, FOXP3+ |
| **NK cells** | Singlet, Live, CD45+, CD3-, NK1.1+ |
| **Myeloid cells** | Singlet, Live, CD45+, CD11b+ |
| **M-MDSCs** | Singlet, Live, CD45+, CD11b+, Ly6C+, Ly6G- |
| **G-MDSCs** | Singlet, Live, CD45+, CD11b+, Ly6C-, Ly6G+ |
| **Macrophages** | Singlet, Live, CD45+, CD11b+, Ly6C-, F4/80+ |
| **Dendritic cells (DCs)** | Singlet, Live, CD45+, CD11c+ |
| **cDC1 cells** | Singlet, Live, CD45+, CD11c+, MHCII+, CD103<sup>hi</sup>, CD11b<sup>lo</sup> |
| **cDC2 cells** | Singlet, Live, CD45+, CD11c+, MHCII+, CD103<sup>lo</sup>, CD11b<sup>hi</sup> |

### Serum isolation

Blood was collected by cardiac puncture immediately following animal sacrifice by rising level of CO_2_. Blood was allowed to coagulate for 30 minutes and then centrifuged at 12,000 x g for 15 minutes at 4°C. Serum was removed from the separated pellet of coagulated blood and stored at -80°C.

### Untargeted metabolomics of tumour samples

The untargeted metabolomics assay was performed by Biocrates (Innsbruck, Austria). The commercially available MxP® Quant 500 kit from Biocrates was used for the quantification of endogenous metabolites of various biochemical classes. Lipids and hexoses were measured by flow injection analysis-tandem mass spectrometry (FIA-MS/ MS) using a SCIEX API 5500 QTRAP® (AB SCIEX, Darmstadt, Germany) instrument with an electrospray ionization (ESI) source, and small molecules were measured by liquid chromatography-tandem mass spectrometry (LC MS/MS), also using a SCIEX API 5500 QTRAP@ (AB SCIEX, Darmstadt, Germany) instrument. The experimental metabolomics measurement technique is described in detail by patents EP1897014B1 and EP1875401B1. Briefly, a 96-well based sample preparation device was used to quantitatively analyse the metabolite profile in the samples. This device consists of inserts that have been impregnated with internal standards, and a predefined sample amount was added to the inserts. Next, a phenyl isothiocyanate (PITC) solution was added to derivatise some of the analytes (e.g., amino acids), and after the derivatization was completed, the target analytes were extracted with an organic solvent, followed by a dilution step. The obtained extracts were then analysed by FIA-MS/ MS and CC-MS/ MS methods using multiple reaction monitoring (MRM) to detect the analytes. Concentrations were calculated using appropriate mass spectrometry software (Sciex Analyst®) and data were imported into biocrates’ MetIDQTM software for further analysis.

Metabolomics analysis to generate PCoA plots, heatmaps and differential metabolite comparisons were conducted using the MetaboAnalyst (V5.0) platform. The data was first normalised by sum, log_10_ transformed, and auto scaled prior to univariate analyses, as described previously (*27*). Differential metabolite levels were compared by two-sample t test with statistical FDR threshold value < 0.05. Metabolite abundances were then input into the MetaboAnalayst platform to allow comparisons of perturbed metabolic pathways, as previously described (*28*).

### Mesoscale discovery (MSD) cytokine analysis of tumour samples

Tissue samples were weighed into a MPBio Lysing Matrix E bead beating tube (MPBio) with 1ml of homogenisation buffer (150 mmol/L NaCl, 20 mmol/L Tris, 1 mmol/L EDTA, 1 mmol/L EGTA, 1% Triton X-100, pH 7.5+ cOmpleteTM protease inhibitor (Roche)). An MPBio Fast Prep bead beater (MPBio) was used to homogenise the tissues at a speed 4.0 for 40 seconds, followed by speed 6.0 for 40 seconds. Samples were centrifuged at 12,000 x g for 12 minutes (4°C) and then stored at −80°C until analysed. Samples were run on a custom Mesoscale Discovery U-PLEX Mouse Kit (MSD) according to the manufacturer’s instructions. Plate was read using an MSD QuickPlex SQ 120 imager (MSD, Rockville, MD, USA).

### PyMT tumour microbiome analysis

Mouse mammary tumours from the indicated anatomical regions were dissected in sterile conditions in tissue culture hoods with autoclaved tools, reagents, and protective equipment. Tumour samples were manually homogenised in 1ml of PBS, with 100μL of the resulting homogenate spread on microbial culture plates. For aerobic culture, homogenate was spread on Columbia blood agar (CBA) (Oxoid) + 5% horse blood; Man Rogosa Sharpe (MRS) agar (Oxoid); brain heart infusion (BHI) agar (Oxoid), and incubated aerobically at 37°C for five days. For anaerobic culture, homogenate was spread on MRS agar (+ 50mg/L-cysteine), BHI agar (+ 50mg/L-cysteine), Peptone Yeast Extract Glucose Starch (PYGS) agar (Thermo) and cultured anaerobically at 37°C for three days. Dissection tools were dipped in sterile PBS and plated on under the same conditions as an environmental control, whilst mouse skin swabs were plated as a positive control to confirm culture conditions could support microbial growth.

### Bifidobacterium pseudocatenulatum qPCR

Bacterial load of *B. pseudocatenulatum* 210 was estimated by qPCR using a species-specific primer (GroEL gene) designed previously by Junick and Blaut (2012) (*29*). The reactions were performed in duplicate in 12.5μl of LightCycler® 480 SYBR Green I Master (Roche Diagnostics, cat. 0470751600), 2.5μl of forward and reverse 10μM primers, 7.3μl of RNase-free water (Qiagen), and 0.2μl of template DNA. Standard curves were generated by serial dilutions of 100 ng/μL of monoculture-extracted DNA to reach the lowest concentration of 0.001 ng/μL. Samples were run on the LightCycler® 480 system (Roche) with the programme as follows: 5 minutes incubation at 95°C, followed by 45 cycles with 15 seconds at 94°C, 15 seconds at 64°C and 15 seconds at 72°C. The melting curve analysis followed with 5 seconds at 95°C, 5 minutes at 65°C and continuous temperature increase to 97°C. Samples were finally cooled to 40°C for 30 seconds before completion. Data was analysed with the LightCycler® 480 Software (v.1.5) (Roche).

### Exopolysaccharide isolation and purification

EPS isolation and purification were performed based on the protocol by Ruas-Madiedo (2021) (*30*). A 20ml bacterial suspension in MRS broth was cultured overnight at 37°C under anaerobic conditions. 200μl of the liquid culture were plated on MRS + L-cysteine (50mg/L), MRS + L-cysteine (50mg/L) + fructose, or MRS + L-cysteine (50mg/L) + arabinose agar plates and incubated for 96 hours anaerobically at 37°C. The bacterial lawn was harvested by adding 1ml of MilliQ water to the plate and scraping with a spreader. This step was repeated as required. One volume of 2M NaOH was added to the harvested bacterial biomass and the resulting solution was stirred gently for 16 h at 150rpm. Next, the bacterial biomass solution was centrifuged for 25 minutes at 9200 rpm at 4°C and the resulting supernatant was collected. EPS was precipitated by adding two volumes of ice-cold absolute ethanol to the supernatant and storing at 4°C for at least 48 hours. The precipitated mixture was centrifuged for 25 minutes at 9200 rpm at 4°C. The supernatant was discarded, the precipitate was dissolved in 10ml MilliQ water and transferred to a pre-soaked Spectra/Por® Dialysis Membrane (Spectrum Laboratories, Inc., USA) with a molecular weight cut off of 8,000 or 10,000 Da. The dissolved precipitate was dialysed against MilliQ water with a daily water change for at least 48 hours. The dialysis product was transferred to sterile empty petri dishes (5ml/petri dish), covered with parafilm which was then poked, and lyophilized overnight using Alpha 1-4; CHRIST LOC-1m (Christ, Germany) lyophilizer. The lyophilized crude EPS (cEPS) was harvested using a 10μl sterile plastic loop, transferred to a sterile cryotube and stored at 4°C.

20mg of crude EPS were dissolved in 4ml of Buffer I (50 mM Tris-HCl pH 7.5, 10 mM MgSO4*7H2O), after which 1000x stock of DNase I (dissolved in Buffer I) was added to a final concentration of 5.5μg/ml. The solution was gently stirred on a shaker at 37°C for 6 hours. Then, stock of 100x Pronase E dissolved in Buffer II (50 mM Tris-HCl pH 7.5, 2% EDTA pH 7-8) was added to a final concentration of 50μg/ml. The solution was gently stirred on a shaker for 18 hours at 37°C. As a next step, 60% TCA was added to a final concentration of 12%. The solution was transferred to 2ml Eppendorf tubes and incubated on a shaker at 21.5°C for 30 minutes at 350 rpm, after which it was centrifuged at 13,000 x g for 25 minutes at 4°C. The supernatant was collected and adjusted to a pH of 5 with 10M NaOH. The solution was transferred to a pre-soaked Spectra/Por® Dialysis Membrane (Spectrum Laboratories, Inc., USA) with a molecular weight cut off of 10,000 Da and dialysed against MilliQ H_2_O at 4°C for 48 hours with a daily water exchange. The dialysed sample was then lyophilised and transferred to a sterile cryotube and stored at 4°C.

### Statistical analysis

Statistical analyses were performed using GraphPad Prism 9 software. Unless otherwise stated, Kolgorov-Smirnov tests were performed to confirm normality of data and Student’s t-test (unpaired, two-tailed, at 95% confidence interval) were used to generate P-values. Where multiple t-tests were performed, a false discovery rate (FDR) of q<0.05 was used. Significant observations are represented according to the following annotation: ****P < 0.001, ***P < 0.001, **P < 0.01, *P < 0.05. In some figures, raw P-values which did not meet statistical significance are presented next to the corresponding data. Where no P-value is presented, statistical analyses did not identify a significant observation. Full details of specific statistical tests performed corresponding to each dataset can be found in the respective figure legends. Quantifications show mean values ± SEM unless otherwise indicated.

## Results

### *Bifidobacterium* cocktail treatment reduces tumour burden in breast cancer models

To explore the impact of *Bifidobacterium* species and strains in breast cancer treatment, we orally treated animals bearing BRPKp110 luminal A-like breast tumours (*31, 32*) with a *Bifidobacterium* cocktail (*Bif*-cocktail), comprised of four distinct strains: *B. bifidum* LH80, *B. pseudocatenulatum* 210, *B. reuteri* LH506, and *B. longum subsp. longum* NCIMB 8809 (Figure 1A). *Bif*-cocktail treatment led to a significant reduction in BRPKp110 tumour volume (Figure 1B) and reduced early dissemination of GFP^+^ tumour cells to the lungs (Figure 1C-D). To dissect the microbial mechanisms driving tumour response, we deconstructed the *Bif*-cocktail, testing the combined treatment against each constituent strain individually in the luminal B-like PyMT-BO1 breast tumour model (*33*). Whilst the *Bif*-cocktail was ineffective in this model, three of the constituent strains (LH80, 210, and LH506) induced an anti-tumour response (Figure 1E). Comparable tumour reductions were observed after administration of the effective single strains in the BRPKp110 model (Figure 1F), albeit with slightly less potent efficacy with strain LH506. The activity of CD8^+^ T cells is consistently associated with the anti-tumour activity of *Bifidobacterium* (*4, 10, 15*) and is a vital pathway mediating patient outcomes. Only strain 210 treatment induced CD8^+^ polarisation to an effector-memory subtype in PyMT-BO1 tumours (Figure 1G), suggesting a CD8^+^T cell-dependent mechanism driving 210 efficacy, and alternative mechanisms mediating response to LH80 and LH506.

**Figure 1.**
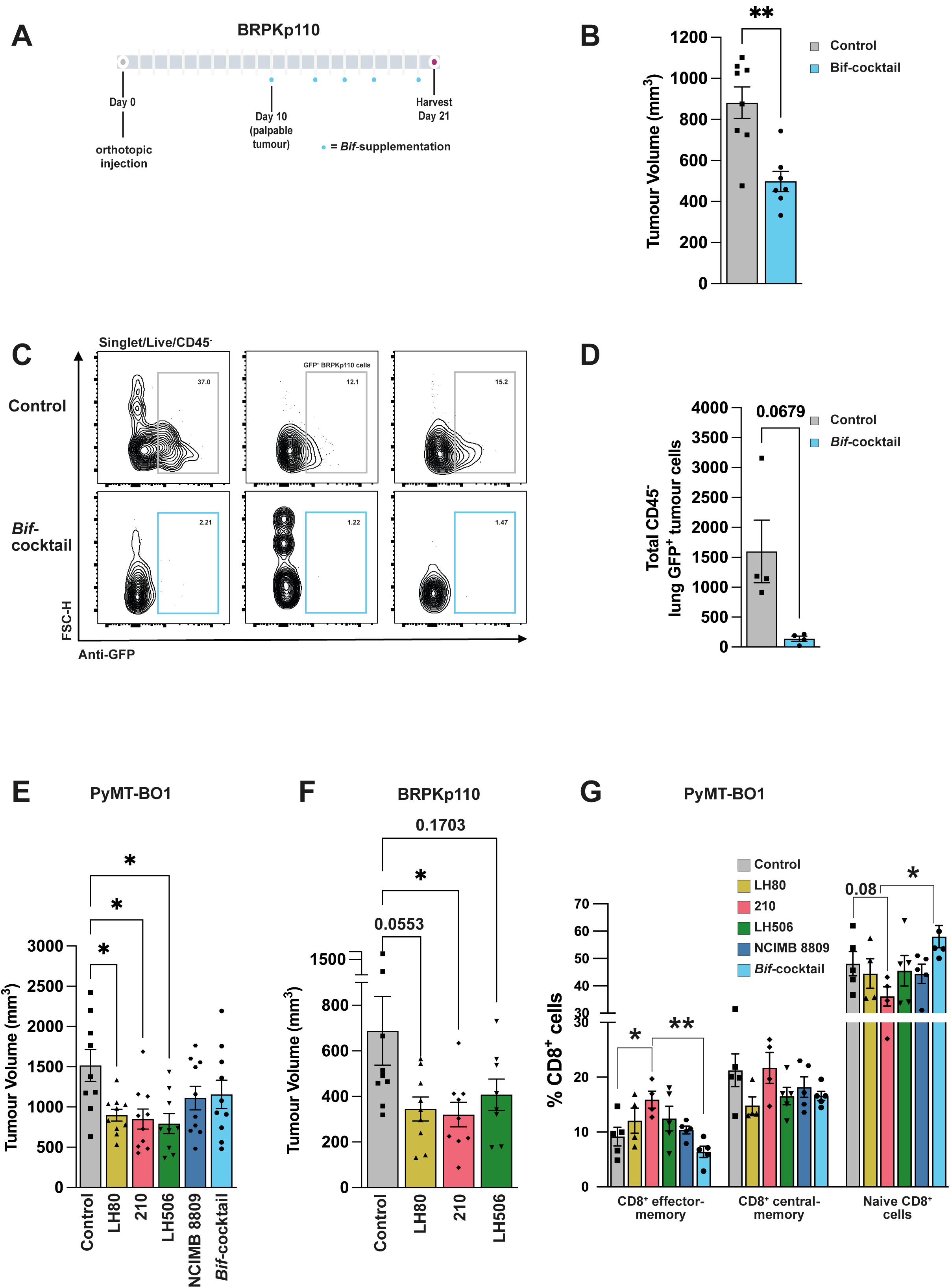
*Bifidobacterium* treatments induce anti-tumour efficacy in mouse breast models. (A) Experimental design BRPKp110 tumour growth experiments. C57 BL/6 mice were orally dosed with ∼1×10^10^ CFU/ml *Bifidobacterium* thrice weekly upon the onset of a palpable tumour (day 10). (B) Primary tumour size of Bif cocktail-treated animals compared with PBS vehicle control. n=7-8. (C) Representative flow cytometry plots of treatment and control groups for the assessment of GFP^+^ BRPKp110 cell infiltration into the lungs of primary tumour-bearing animals. (D) Quantification of GFP^+^ BRPKp110 cell infiltration to the lungs following treatment with the Bif cocktail, n=4. (E) Endpoint PyMT-BO1 tumour volumes following administration of various unique strains of Bifidobacterium, or the four-strain consortia (Bif cocktail) comprised of each the individual strains. n=9-10. (F) Endpoint BRPKp110 tumour volumes following administration of *Bifidobacterium* strains. n=7-9. (G) Quantification of CD8^+^ effector-memory polarisation following administration of *Bifidobacterium* treatments. n=4-5. Statistical significance calculated by (B and D) two-tailed unpaired t test or (E-G) one-way ANOVA with Tukey’s multiple comparisons test. **P < 0.01, *P < 0.05.

### *B. pseudocatenulatum* 210 exhibits consistent anti-tumour activity across Breast Cancer models and enhances standard-of-care therapy

We focused on *B. pseudocatenulatum* 210 due to its strong and consistent anti-tumour effects. When administered as a monotherapy, 210 induced consistent anti-tumour efficacy across several pre-clinical orthotopic models representing luminal A (BRPKp110), luminal B (PyMT-BO1), and triple negative (4T1) breast cancer – significantly reducing tumour burden (Figure 2A). In addition, histological analyses showed that 210 also reduced 4T1 metastatic spread to the lungs (Figure 2B-C), demonstrating its systemic anti-tumour efficacy. In the autochthonous PyMT model, 210 monotherapy not only reduced tumour burden, but also delayed tumour formation, suggesting its potential for longer-term anti-tumour activity and prophylactic utility (Figure 2D). Furthermore, administration of 210 enhanced the efficacy of standard-of-care treatments, including cyclophosphamide chemotherapy in luminal BRPKp110 tumours and anti-PD-1 immunotherapy in triple negative 4T1 tumours (Figure 2E-F).

**Figure 2.**
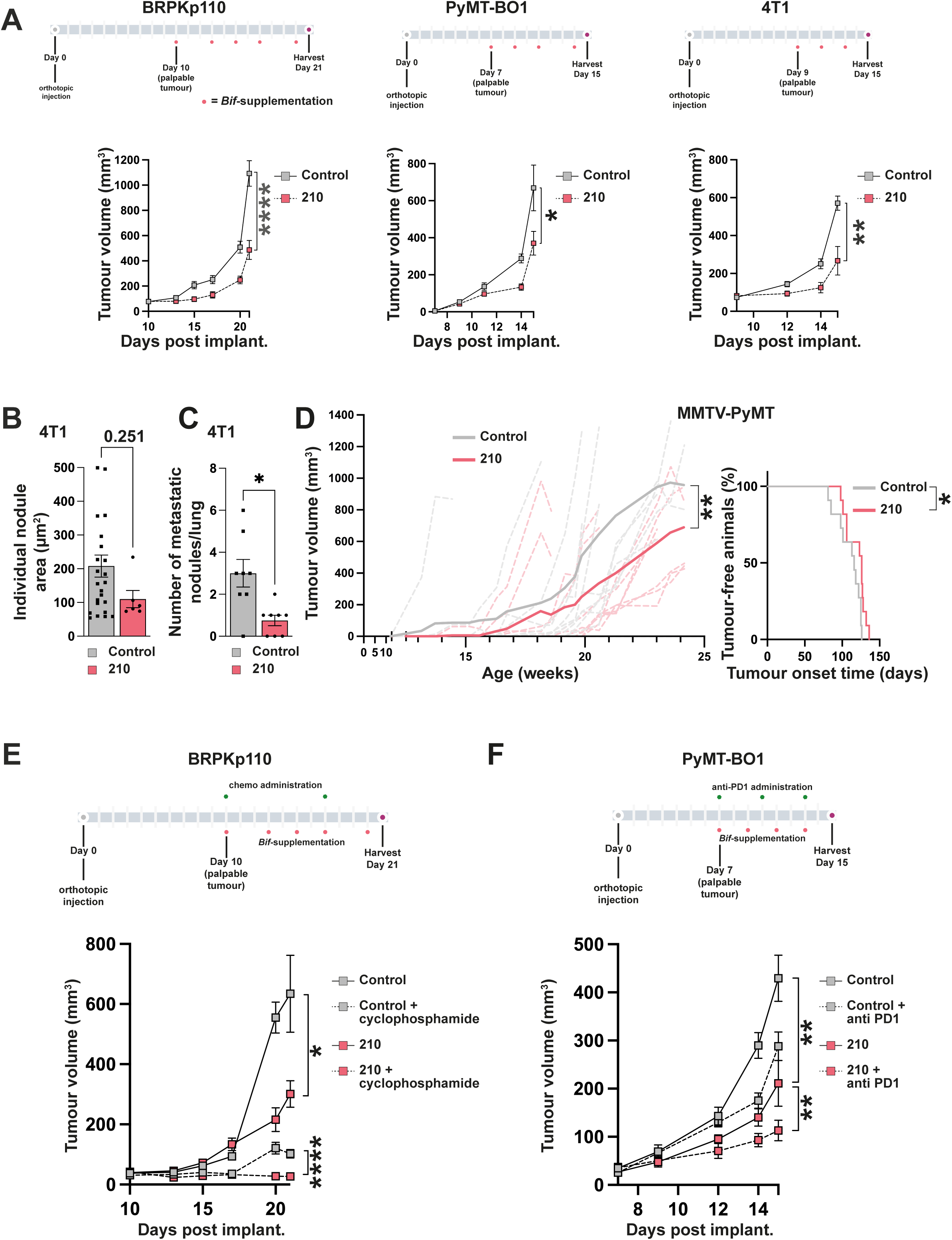
*B. pseudocatenulatum* 210 treatment inhibits breast tumour progression and enhances response to standard of care therapies. (A) Experimental outlines and tumour growth responses to 210 treatment in the BRPKp110 (n=17-18, N=2), PyMT-BO1 (n=8-9), and 4T1 (n=9) orthotopic tumour models. (B) Quantification showing the average size (n=6-24) and (C) number (n=9) of macromolecular metastatic nodules observable in the lungs of 4T1-bearing animals visualised following histological H&E staining. (D) MMTV-PyMT spontaneous tumour burden growth and tumour onset time following 210 administration. n=12. (E) Experimental outline and associated BRPKp110 tumour growth responses of animals administered combinations of 210 and cyclophosphamide chemotherapy (n=6-8), and (F) 4T1 tumour-bearing animals treated with combinations of 210 and anti PD-1 immunotherapy. Statistical significance calculated by (A, D, E, F) two-tailed unpaired t test, (B) Mann-Whitney test, (C) two-tailed unpaired t test with Welch’s correction, and (D) Kaplan-Mieier survival analysis of tumour free animals. ****P < 0.0001, **P < 0.01, *P < 0.05.

### CD8+ T cells drive the anti-tumour response to *B. pseudocatenulatum* 210

Next, we wanted to characterise the immune response driving tumour inhibition of 210. No significant changes were observed in the gross infiltration of major lymphoid populations in the tumour microenvironment (Figure S1A), suggesting immune effects were more likely caused by differential polarisation and activation of immune populations. In agreement with our initial data in PyMT-BO1 tumours, 210 administration also increased BRPKp110 CD8^+^ T cell polarisation towards an effector-memory subtype (Figure 3A), albeit not to a statistically significant level in the BRPKp110 model. CD8^+^ cells in the spleen and tumour-draining lymph node (tdLN), were more polarised to the central-memory subtype, a state associated with long-lived antigenic memory (Figure 3B). Additionally, blood circulating CD8^+^ T cells showed enhanced activation, marked by increased expression of the CD44 activation marker (Figure 3C). Cytokine analysis demonstrated that 210 induced increases in TNFa and (non-statistically significantly) IFNy in the primary tumour (Figure 3D) and were mirrored by a significant increase in serum IFNy levels (Figure 3E). BRPKp110 tumour infiltrating CD8^+^ cells in both BRPKp110 and 4T1 tumour models (Figure 3F-G), secreted higher levels of TNFa and IFNy following 210 administration suggesting their involvement in gross increases observed for these cytokines. Enhanced cytolytic activity of CD8^+^ T cells from 210-treated animals was confirmed by increased granzyme B and CD107a expression (Figure 3H-I). Concurrently, tumour infiltrating T helper cells were not more polarised or activated, aside from a small increase in IL-4 production restricted to the BRPKp110 model (Figure S1B-E). Likewise, BRPKp110 tumour-infiltrating NK cells were not more abundant and did not produce any higher levels of inflammatory cytokines (including IFNy and TNFa) (Figure S1F-G), suggesting CD8^+^ T cells may be the sole cytotoxic effector population mediating 210 anti-tumour response. Notably, depletion of CD8^+^ T cells *in vivo* in BRPKp110 tumour-bearing animals abolished the anti-tumour efficacy of 210, validating a CD8^+^-dependent mechanism of action (Figure 3J).

**Figure 3.**
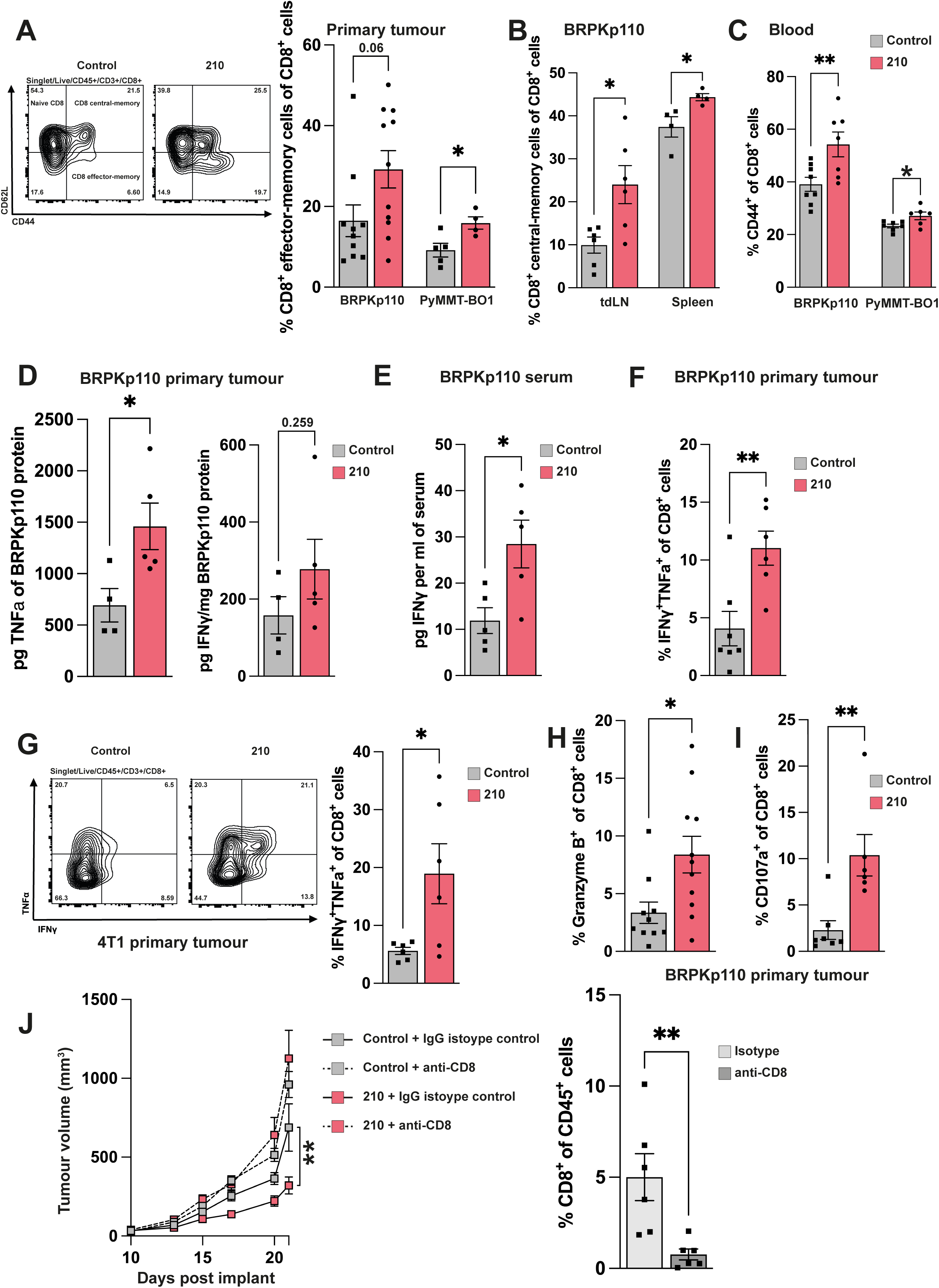
*B. pseudocatenulatum* 210 anti-tumour efficacy is dependent on CD8^+^ T cell activity. (A) Representative flow cytometry plot showing with quantification showing effector-memory CD8^+^ T cell polarisation in PyMT-BO1 primary tumours following 210 administration. n=11-12, n=5. (B) Quantification showing CD8^+^ T cell central-memory cell polarisation (n=6, n=4) and (C) CD44^+^ expression (n=8, n=7) in the indicated tumour models and tissues. (D) Quantification of IFNγ and TNFα levels (n=4-5) in primary tumours and (E) IFNγ levels in the sera (n=5) of BRPKp110-bearing animals measured by MSD multiplex cytokine analysis. (F) Co-expression of IFNγ and TNFα (n=6-7) by BRPKp110 primary tumour CD8^+^ T cells following 210 treatment. (G) Representative flow cytometry plot with quantification of IFNγ and TNFα co-expression by 4T1 primary tumour CD8^+^ T cells following 210 treatment. n=6. (H) Expression of granzyme B (n=11-12), (I) CD107a (n=6-7) by BRPKp110 primary tumour CD8+ T cells following 210 treatment. (J) BRPKp110 mean tumour growth over time following administration of vehicle control or 210 in combination with either anti CD8-depleting antibody or IgG isotype control (n=9), with quantification of the depletion of primary tumour CD8^+^ T cells following antibody administration. Statistical significance calculated by (A-F, H-J) two-tailed unpaired t test and (G) Mann-Whitney test. **P < 0.01, *P < 0.05.

### *B. pseudocatenulatum* 210 alters tumour-associated macrophages and enhances DC activation

Alongside the CD8^+^ T cell response, we investigated the activity of innate immune cells given they orchestrate T cell populations. Within BRPKp110 and PyMT-BO1 primary tumours, we observed a significant reduction in CD206^+^ tumour-associated macrophages (TAMs) (Figure 4A-D). These CD206^+^ TAMs, historically characterised as ‘M2-like’ and pro-tumourigenic (*34*), were replaced with, MHCII+ TAMs (‘M1-like’) indicative of a shift towards a more anti-tumourigenic, CD8^+^-privileged tumour microenvironment (TME) (Figure 4B). Although the inflammatory state of the TME is vital, we also showed systemic activation of CD8^+^ T cells (in the spleen, tdLN, and blood), which may reflect upstream pathway activation away from the localised TME. Systemic analysis of circulating innate cells highlighted a near-significant increase in the circulating pool of DCs (Figure 4E). DC infiltration and activation are pivotal for priming and activation of CD8^+^ T cells due to their role as antigen presenting cells (*35, 36*). Systemic analysis indicated that 210 induced increases in circulating DCs; particularly in the CD8^+^-specific cDC1 population, while no significant changes were observed in the T helper-specific cDC2 population (Figure 4F). This suggests that 210 may preferentially stimulate CD8+ T cell responses via cDC1, pointing to a potential mechanistic cascade for its anti-tumour efficacy. Increased cDC1 levels were also observed within the tdLN (Figure 4G), a key site for anti-tumour CD8^+^ programming. Additionally, DCs in the tdLN of 210-treated animals showed elevated expression of CD80^+^CD86^+^ maturation markers, approaching statistical significance (Figure 4H). This suggests that 210 enhances the maturation of DCs, potentially boosting their ability to cross-present tumour-associated antigens and effectively stimulate naïve CD8+ T cells towards an effector phenotype. To further assess whether 210-induced DC activation was directly responsible for the enhanced anti-tumourigenic efficacy and increased CD8^+^ activity, we co-cultured BMDCs with 210 cells and adoptively transferred the treated BMDCs to naïve animals bearing BRPKp110 tumours (Figure 4I). Notably, 210-conditioned BMDC treatment significantly reduced BRPKp110 tumour burden and enhanced CD8^+^ cell activity within the TME, as evidenced by increased production of inflammatory IFNy cytokine. This confirms that 210-activated DCs are capable of robustly stimulating anti-tumour CD8^+^ T cell immunity (Figure 4J).

**Figure 4.**
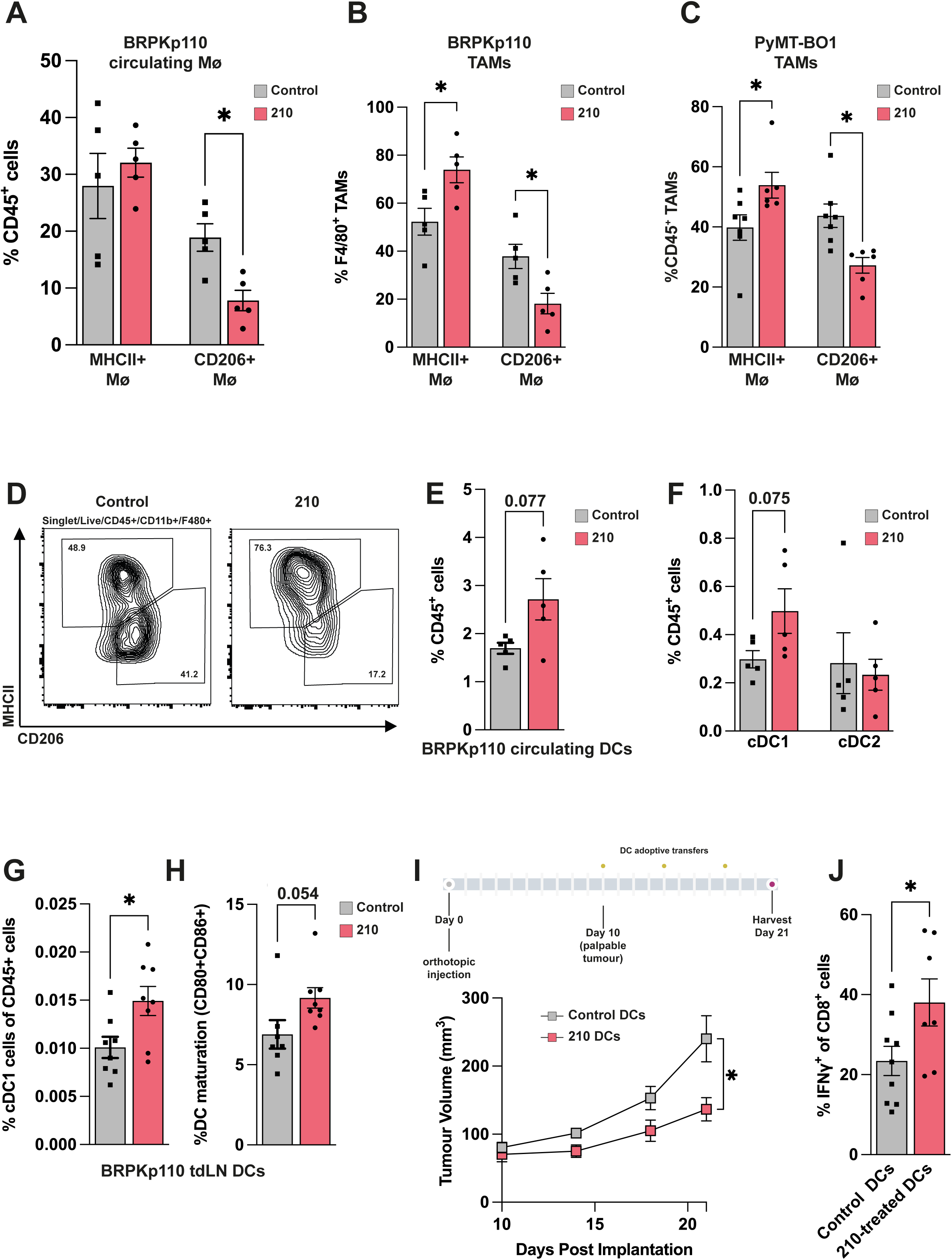
*B. pseudocatenulatum* 210 programmed dendritic cells induce anti-tumour CD8^+^ T cell immunity. (A) Quantification of the infiltration and (B) polarisation of MHCII^+^ and CD206^+^ macrophages within BRPKp110 (n=5) and (C) PyMT-BO1 (n=6-7) primary tumours following 210 administration, with (D) a representative flow cytometry plot from BRPKp110 tumours. (E) Quantification of the infiltration of dendritic cells and (F) cDC subtype cells in the blood of BRPKp110-bearing animals (n=5). (G) Quantification of the infiltration of cDC1 cells (n=8) and (H) the percentage of CD80^+^CD86^+^ mature dendritic cells (n=7-8) within BRPKp110 tumour-draining lymph nodes following 210 administration. (I) Experimental outline of 210-treated dendritic cell adoptive transfers to naïve BRPKp110 tumour bearing animals, with quantification of tumour growth (n=8-10) and (J) intratumoural CD8^+^ T cell IFNγ expression (n=7-9) following the indicated treatments. Statistical significance was calculated by two-tailed unpaired *t* test, with Welch’s correction applied to (E) and (F). *P < 0.05.

### *B. pseudocatenulatum* 210 anti-tumour effects are mediated by EPS

Having defined a DC and CD8^+^ T cell-based mechanism underpinning 210 anti-tumour efficacy we next wanted to identify the functional compound(s) driving these host-microbe interactions. While immunogenic metabolite secretion is a commonly described mechanism in bacterial-mediated cancer immunity (*37*) untargeted metabolomics analysis of the serum from BRPKp110 and PyMT-BO1 tumour-bearing animals did not identify any significant changes to the global metabolome profile, or to the abundance of individual circulating metabolites following 210 administration (Figure 5A-B). Furthermore, differential expression analysis of serum metabolites across both models showed no consistent patterns of change (Figure 5C), suggesting that circulating metabolites were unlikely to be responsible for the observed 210-induced anti-tumour effects. To gain further clarity on the microbial dynamics and activity of 210, we conducted colonisation studies in germ free monocolonised mice, using standard oral administration at 1×10^10^ CFU. As expected, no viable bacteria were detected in vehicle control mice, whilst 210 monocolonised mice only had minimal viable bacterial colonies, localised primarily in the colon and caecum (Figure S2A). Interestingly, live 210 cells were not able to establish prolonged colonisation and were largely cleared from the gut within 24 hours post-administration. This suggested a transient interaction between 210 and the host. Further validation using *B. pseudocatenulatum* specific PCR analysis of wild-type animals, confirmed that 210 cell load was most prominent in the caecum and colon, with 210 signature detectable up to 24 hours post-administration (Figure S2B). Despite the clearance of live bacteria, 210-specific signatures (GroEL gene), were detectable in faecal pellets. 210 gene signatures returned to approximate baseline levels up to 48 hours post oral administration, indicating that dead bacterial cells or bacterial fragments may persist longer in the gut environment (Figure S2C). This persistence suggests that even non-viable bacterial components may play a role in modulating the immune response. Next, we sought to determine whether 210 could translocate to or colonise tumour tissues. Selective qPCR of BRPKp110 primary tumours (Figure S2D) and culturomic analysis of MMTV-PyMT primary tumours (Figure S2E) failed to detect any 210 cells, suggesting the absence of a direct tumour microbiome-based mechanism. To further investigate if 210’s anti-tumour effect was dependent on other gut commensal bacteria, we pre-treated BRPKp110 tumour bearing animals with a broad-spectrum antibiotic cocktail to deplete the native microbiota. Notably, antibiotic treatment did not rescue tumour growth inhibition in 210-treated animals, suggesting 210 exerts it effects independently of the commensal microbiome and directly induces anti-tumour immunity (Figure S2F).

**Figure 5.**
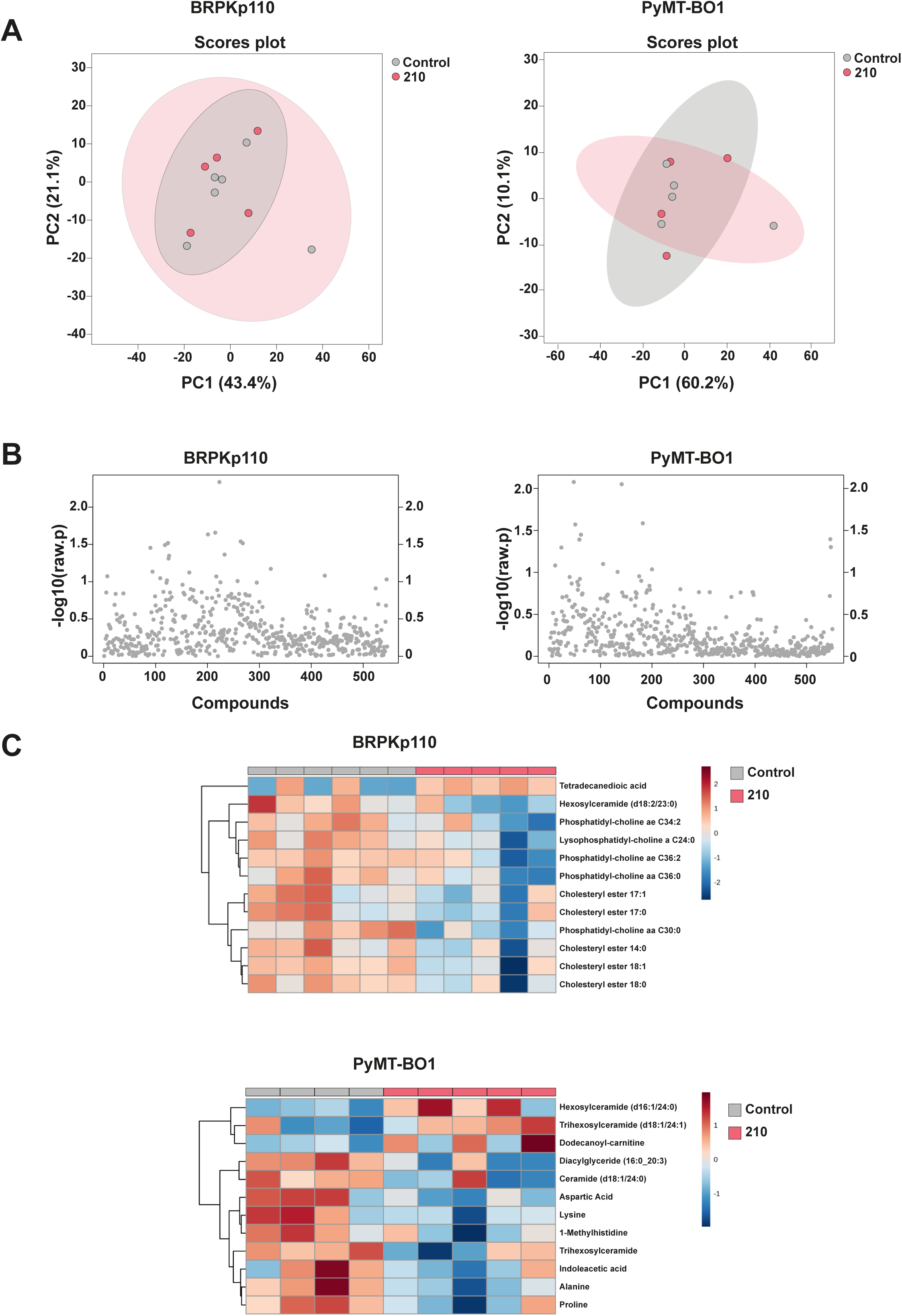
Administration *of B. pseudocatenulatum* 210 does not significantly alter serum metabolite levels in untargeted analyses. (A) 2D score plots of serum metabolite profiles of BRPKp110 (n=5-6) and PyMT-BO1 (n=4-5) tumour-bearing animals; shaded circles indicate 95% confidence intervals. (B) Plots showing significance scores for the differential expression of 500 serum metabolites following 210 therapeutic intervention, n=5. (C) Heatmaps showing the top 12 differentially expressed serum metabolites between vehicle control and *B. pseudocatenulatum* 210 treated animals. Statistical differences were assessed by two-tailed unpaired *t* test with an FDR applied at *P* < 0.05

The absence of involvement from other commensals, combined with the poor colonisation and viability of 210, and lack of 210-induced metabolic changes, suggested a mechanism driven by bioactive structural components rather than active microbial processes. To test whether viability of 210 cells was necessary for therapeutic efficacy, we administered peracetic acid-killed 210 cells, a method shown by *Moor et al* (*26*) to effectively kill microbes whilst preserving their surface structures. Acid killed 210 was able to inhibit BRPKp110 tumour growth *in vivo* to a similar extent as live 210 cells (Figure 6A), confirming that 210’s anti-tumour activity is not dependent on metabolite production or live cell interactions. Given that *Bifidobacterium* species and strains are known to produce immunogenic EPS on their cell surfaces, which have been shown to mediate specific immune responses in lymphoid populations (*38*) and DCs (*23*), we hypothesized that 210 EPS might be the key driver of its anti-tumour effects. To evaluate this, we administered isolated preparations of 210 EPS to BRPKp110 tumour bearing animals. Strikingly, 210 EPS treatment significantly reduced tumour progression, mirroring efficacy of live 210 cells (Figure 6B). Furthermore, 210 EPS replicated the observed increases in intratumoural CD8^+^ T cell cytokine expression and effector-memory differentiation (Figure 6C), suggesting that EPS alone can trigger the same immunological responses as live 210. Within the TME, 210 EPS also reduced the infiltration of CD206^+^ TAMs (Figure 6D). Consistent with live 210 treatment, 210 EPS elevated systemic IFNy levels (Figure 6E), and DC and cDC1 infiltration to levels even higher than those observed with live 210 (Figure 6F-G). Accordingly, 210 EPS did not induce significant changes in infiltration or activity of other lymphoid effector populations, such as T helper or NK cells in the TME (Figure S3A-C). To validate that the 210 EPS-specific interactions with DCs were causative in the mechanism, we performed a BMDC adoptive transfer experiment. BMDCs conditioned with 210 EPS, but not vehicle control or EPS from a *B. longum* strain (B71) previously shown to be ineffective in tumour reduction, were able to induce significant anti-tumour activity in naïve animals (Figure 6H). Moreover, 210 EPS-conditioned BMDCs stimulated robust CD8^+^ T cell activation (Figure 6I) within the tumour. These data demonstrate that *B. pseudocatenulatum* 210 EPS is the key bioactive compound driving the DC-based immune response, leading to CD8+ T cell-mediated tumour suppression.

**Figure 6.**
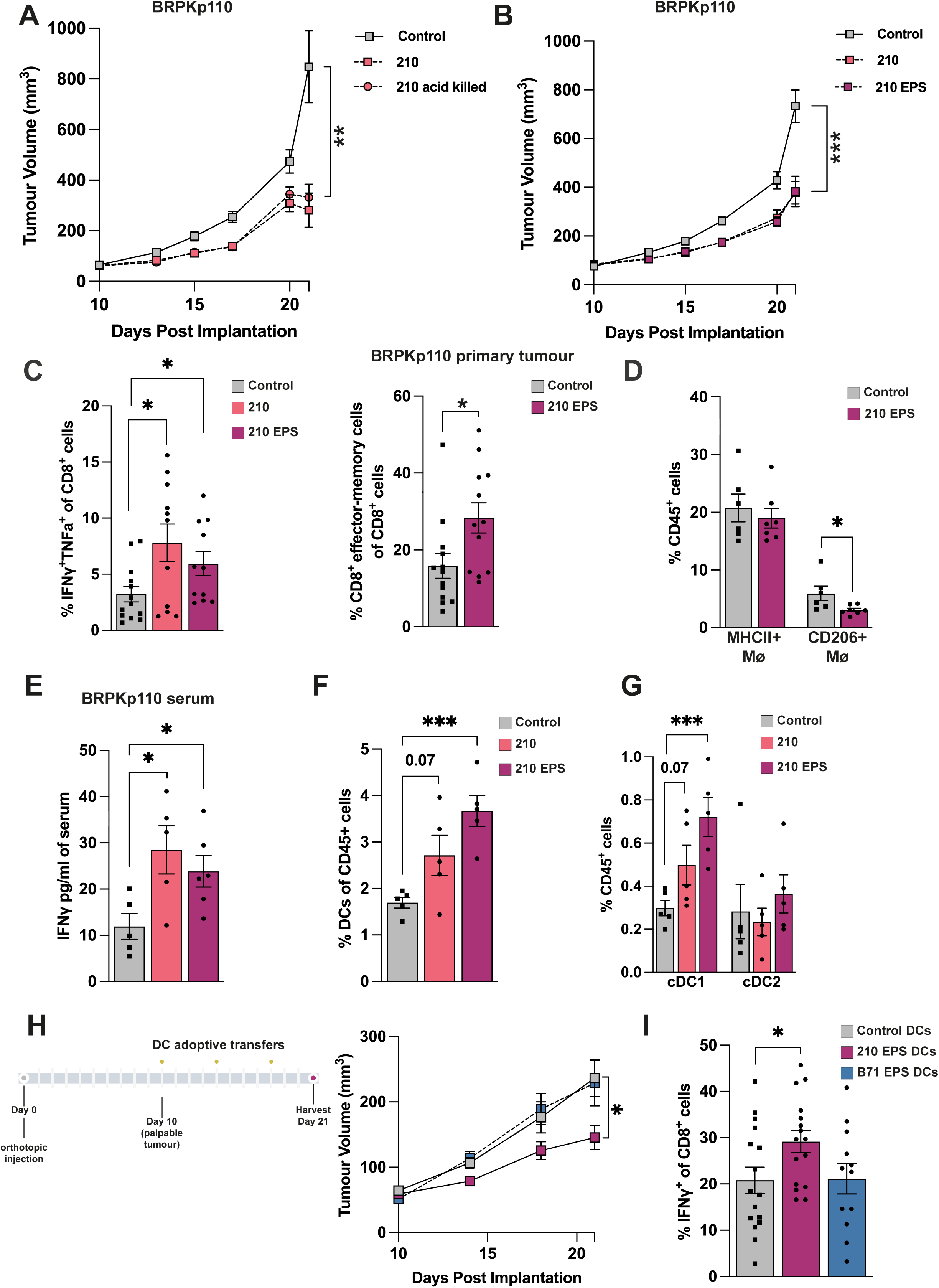
*B. pseudocatenulatum* 210 anti-tumour immunity is induced by cell surface capsular exopolysaccharide. (A) BRPKp110 primary tumour growth following administration with live 210 cells or peracetic acid-killed 210 cells. n=8-9. (B) BRPKp110 primary tumour growth following administration of live 210 or isolated 210 exopolysaccharide (210 EPS) solution (80µg/dose). n=20-23, N=3. (C) Quantification showing BRPKp110 primary tumour CD8^+^ T cell IFNγ and TNFα cytokine expression (n=11-13), effector-memory polarisation (n=11-13), and (D) macrophage MHCII^+^/CD206^+^ infiltration (n=6-7) following administration with 210 EPS. (E) Quantification of IFNγ in the serum of BRPKp110-bearing animals, measured by MSD multiplex cytokine analysis. n=5-6. (F) Quantification of the infiltration of dendritic cells and cDC subtype cells in the blood of BRPKp110-bearing animals. n=5. (G) Experimental outline of B*. pseudocatenulatum* 210 EPS and *B. longum* B71 EPS-treated dendritic cell adoptive transfers to naïve BRPKp110 tumour bearing animals. Data shows quantification of tumour growth (n=18-19, N=2) and intratumoural CD8^+^ T cell IFNγ expression (n=12-16) following the indicated dendritic cell treatments. (A-H) Statistical significance was calculated by two-tailed unpaired *t* test. ***P < 0.001, **P < 0.01, *P < 0.05.

## Discussion

Therapeutically harnessing gut microbes to treat chronic disease is a fundamental aim in microbiome research. Whilst numerous studies have highlighted the potential of microbiota-based interventions, the success of translating these findings to extra-intestinal disease therapies remains limited. A significant barrier to this translation is our incomplete understanding of the specific microbial-derived functional compounds that drive therapeutic responses. Without identifying these key compounds, predicating how they may be influenced by variable patient-specific conditions becomes difficult. In this study, we demonstrate the broad potential of various species of *Bifidobacterium* to treat breast cancer. We provide strong evidence that *B. pseudocatenulatum* 210 inhibits breast tumour progression and enhances the efficacy of standard-of-care therapies through presentation of cell surface capsular EPS. This is the first *in vivo* study demonstrating *Bifidobacterium* EPS as an anti-tumour compound, advancing the concept of microbial polysaccharides as novel immunotherapy agents.

The link between *Bifidobacterium* and cancer has been an active area of research for the past decade. Sivan et al, (*4*) were the first to show that a cocktail of four *Bifidobacterium* species enhanced immune checkpoint inhibitors (ICIs) responsiveness to melanoma, a mechanism dependent on DCs-and live bacteria. While their study implicated secreted metabolites, it left open questions about the specific microbial components driving the response. Subsequent studies have demonstrated the anti-tumour potential of single *Bifidobacterium* strains (*10, 15, 19*), including in colorectal cancer and melanoma models.

However, few studies have explored the efficacy of *Bifidobacterium* in breast cancer or identified the microbe-derived active compounds mediating protective effects. In this study, we observed that a *Bifidobacterium* (Bif)-cocktail induced anti-tumour efficacy in luminal A (BRPKp110) but not luminal B (PyMT-BO1) model. Further dissection of the Bif-cocktail revealed that three of its four component strains (*B. bifidum* LH80, *B. reuteri* LH506, and *B. pseudocatenulatum* 210) independently inhibited tumour progression, whilst *B. longum* NCIMB 8809 did not. Intriguingly, this suggested that *B. longum* NCIMB 8809 may have inhibited or diluted the efficacy of the other strains. This highlights a challenge in using microbial consortia or complex communities of live bacteria - interactions within these mixtures may obscure or counteract the therapeutic potential of individual strains. Our findings underscore the importance of tailored microbiome therapies using carefully selected strains to optimise therapeutic pathways. Focusing on the individually effective single strains, analysis of PyMT-BO1 intratumoural CD8^+^ cells showed *B. pseudocatenulatum* 210 induced effector-memory polarisation, whilst *B. bifidum* LH80 and *B. reuteri* LH506 did not. This finding suggests that although multiple *Bifidobacterium* species and strains may have therapeutic potential against (breast) cancer, the underlying molecular mechanisms mediating responses are likely distinct. Given these differences, future therapeutic strategies could benefit from using tailored combinations of microbes to synergistically target multiple therapeutic pathways to enhance patient responses across different cancer subtypes.

Our investigation into *B. pseudocatenulatum* 210 revealed potent anti-tumour effects across multiple breast cancer models, including long-term efficacy in the spontaneous PyMT model and enhanced responses to standard therapies, such as chemotherapy and ICI treatment in luminal and triple-negative breast cancers, respectively. These results are the first to demonstrate the *in vivo* therapeutic potential of *B. pseudocatenulatum* in cancer. Mechanistically 210’s efficacy was shown to be dependent on CD8^+^ T cells, with 210 administration promoting the maturation of cDC1 DCs and stimulating effector-memory polarisation of CD8+ T cells. Importantly, these effects were observed not only within the tumour, but also at systemic sites such as tdLN, spleen, and blood, pointing to a systemic activation of CD8+-driven anti-tumour immunity. Further analysis of the innate immune compartment showed that 210 treatment increased the number of pro-inflammatory macrophages and cDC1 DCs in the blood and tdLN, supporting CD8+ T cell activation and anti-tumour responses. The observed increases in systemic DC maturation and activation, alongside the reduced infiltration of CD206+ TAMs in the TME, suggest 210 induces a systemic shift towards a pro-inflammatory, anti-tumour immune profile. Our adoptive transfer experiments with 210-conditioned BMDCs confirmed the causative role of 210-activated DCs in driving CD8+ T cell-mediated tumour suppression, providing further evidence for the mechanistic basis of 210’s efficacy.

Despite the compelling evidence of *B. pseudocatenulatum* 210’s therapeutic potential, our data showed that 210 did not robustly colonise the gastrointestinal tract of germ-free mice, nor did it induce significant changes to the metabolome. Rather, acid killed 210 maintained anti-tumour efficacy suggesting structural components, rather than live bacterial activity or metabolic output, mediate responses. This led us to hypothesise that 210’s EPS, rather than its metabolites, was the key effector molecule. EPS are known immunogenic components of many gut bacteria, including *Bifidobacterium*, and have been shown to modulate immune responses through interactions with DCs and other immune cells. In our study, isolated EPS from *B. pseudocatenulatum* 210 replicated the anti-tumour effects of live 210, including the stimulation of CD8+ T cell activity, DC maturation, and TAM repolarisation. Adoptive transfer of BMDCs conditioned with 210-derived EPS further confirmed its role as the critical driver of anti-tumour immunity. Our findings align with emerging evidence of microbial polysaccharides as potent immune modulators in cancer therapy. A recent study demonstrated that soluble β-glucan polysaccharide could enhance anti-tumour responses by activating Dectin-1 receptors on immune cells, thus potentiating CD8+ T cell activity(*39*). Similarly, our study demonstrates that 210’s EPS stimulates cDC1 maturation and primes DCs to activate CD8+ T cells, providing new mechanistic insight into the anti-tumour activity of microbial polysaccharides. These results highlight the potential for EPS-based therapies to serve as stand-alone immunotherapeutics or as adjuvants to enhance the efficacy of existing treatments like ICIs.

The clinical relevance of these findings is underscored by the growing body of evidence linking *Bifidobacterium* species to favourable positive cancer outcomes (*12, 40*). Of specific interest, is the correlation of *B. pseudocatenulatum*, alongside other beneficial species like *Roseburia* spp. and *Akkermansia muciniphila*, with enhanced overall response rates and progression-free survival in several cohorts of melanoma patients receiving ICIs (*11*). By demonstrating that *B. pseudocatenulatum* 210 EPS drives CD8+ T cell-mediated anti-tumour immunity in breast cancer, we provide a strong foundation for future clinical translation of *Bifidobacterium*-based interventions, particularly in high-risk populations where disparities in cancer outcomes persist.

In conclusion, this study identifies *B. pseudocatenulatum* 210 EPS as a novel anti-tumour therapeutic compound. Our findings represent a significant advancement in understanding the functional role of microbial polysaccharides in cancer immunotherapy and provide a compelling case for further exploration of microbial EPS in therapeutic contexts. Future work should focus on translating these findings to human clinical trials, with an emphasis on personalising microbiota-targeted therapies to improve patient outcomes.

## Supplementary Figure Legends

**Figure S1.**
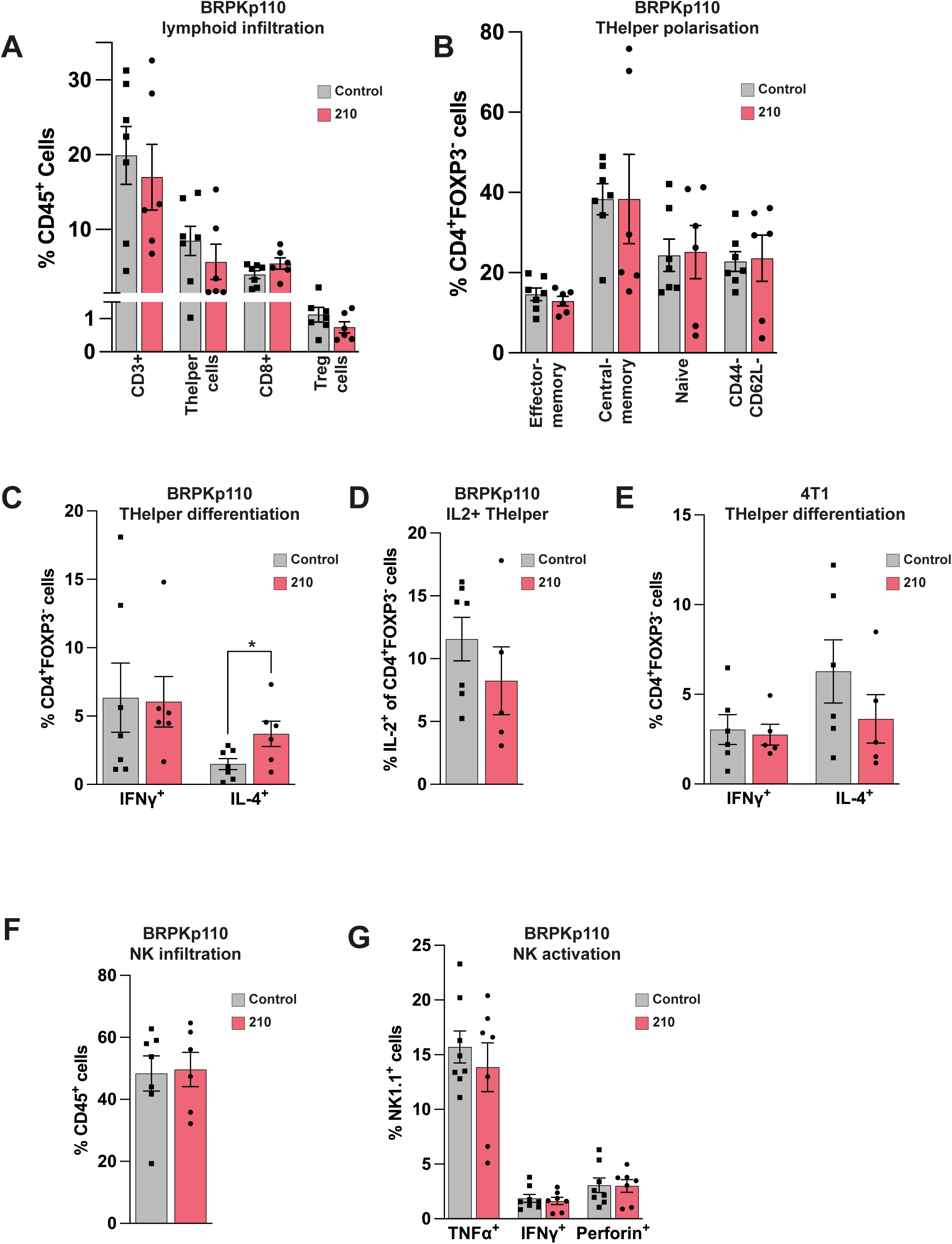
The *B. pseudocatenulatum* 210 immune mechanism does not depend on the infiltration or activation of other lymphoid effector cells. (A) Infiltration of lymphoid immune cell populations into BRPKp110 primary tumours following 210 administration. n=6-7. (B) Primary tumour T helper cell effector-memory differentiation (n=6-7) and (C-D) cytokine expression in BRPKp110 tumours (n=6-7), and T helper cytokine expression in 4T1 tumours following 210 administration (n=5-6). (F) Primary tumour NK cell infiltration (n=6-8) and (G) expression of activation markers TNFα, IFNγ, and perforin (n=7-8). Statistical significance was measured by two-tailed unpaired t test. *P < 0.05.

**Figure S2.**
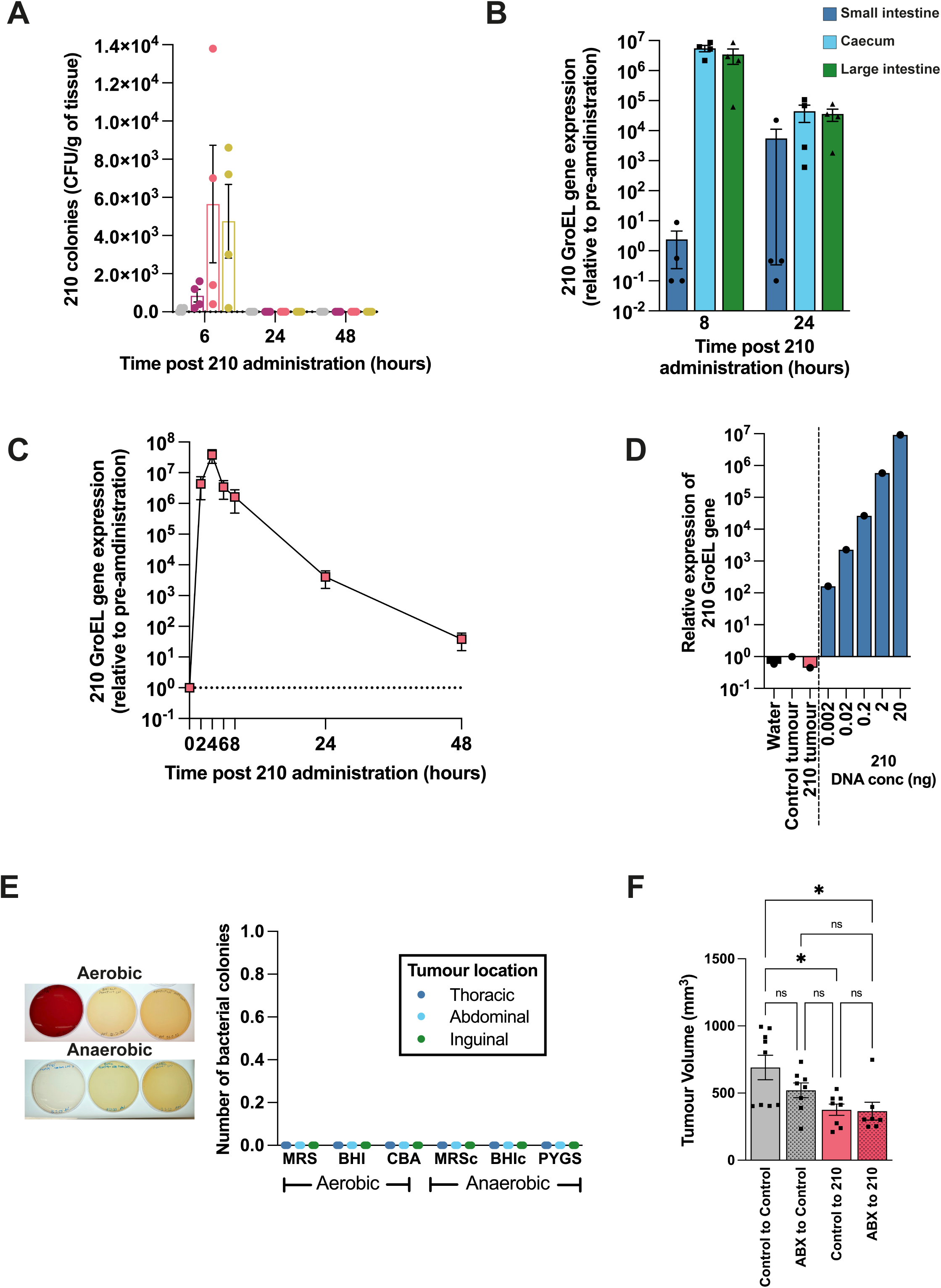
*B. pseudocatenulatum* 210 cells are poorly viable following oral administration and do not modulate the wider gut microbiome. (A) Concentration of culturable colonies of 210 cells isolated from faecal tissue within the gastrointestinal tract of germ free monocolonised animals. n=4. (B) *Bifidobacterium pseudocatenulatum* GroEL gene quantification by qPCR of wildtype gastrointestinal contents, (D) faeces and (E) BRPKp110 primary tumours following 210 administration. n=4. (E) Representative image of agar plates, with quantification of CFU, following culture of homogenised MMTV-PyMT tumour tissues from animals treated with 210. (F) Endpoint BRPKp110 tumour volumes of control or VNMAA antibiotic cocktail pre-treated animals supplemented with either *B. pseudocatenulatum* 210 or vehicle control. n=7-9. Statistical significance was calculated by one-way ANOVA with Tukey’s multiple comparison test. *P < 0.05.

**Figure S3.**
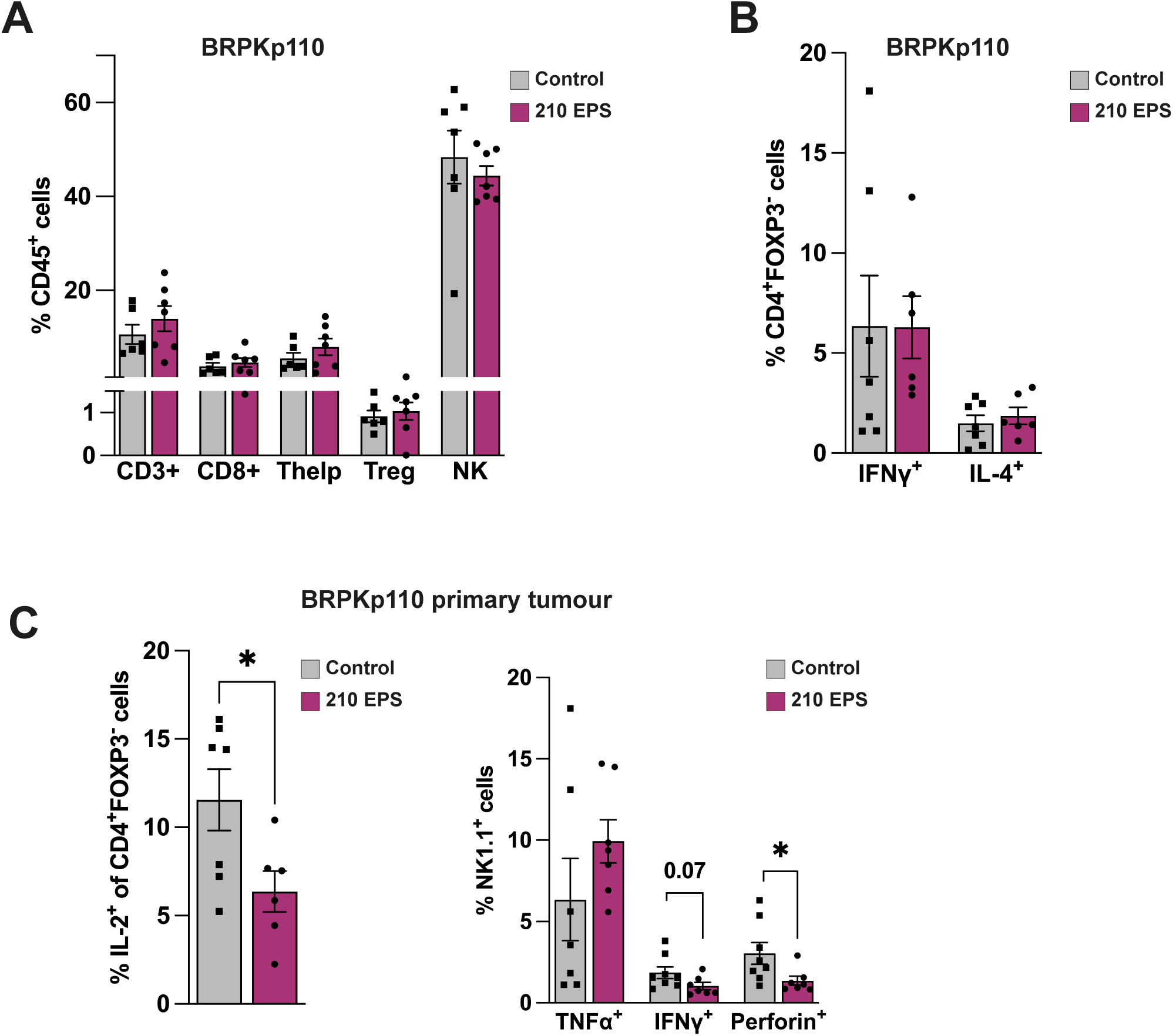
*B. pseudocatenulatum* 210 exopolysaccharide induced anti-tumour immunity is not dependent on other immune effector populations. (A) Infiltration of lymphoid immune cell populations into BRPKp110 primary tumours following 210 EPS administration. n=7. (B) Primary tumour T helper cell IFNγ and IL-4 cytokine expression in BRPKp110 tumours (n=6-7). (C) Primary tumour T helper cell IL-2 expression (n=6-7) and NK cell expression of activation markers TNFα, IFNγ, and perforin (n=7). Statistical significance was measured by two-tailed unpaired *t* test. *P < 0.05.

**Supplementary Table 1.**

| Species | Strain designation | Provenience | Host | Host Gender | Geographical Location |
| --- | --- | --- | --- | --- | --- |
| *Bifidobacterium bifidum* | LH_80 | in-house | Human infant | Male | Norfolk, UK |
| *Bifidobacterium pseudocatenulatum* | 210 | in-house | Human infant | Female | Norfolk, UK |
| *Bifidobacterium reuteri* | LH_506 | in-house | Barbary sheep | n/a | Norfolk, UK |
| *B. longum* subsp. *longum* | NCIMB 8809 | public | Human infant | n/a | n/a |
| B. longum subsp. longum | B_71 | in-house | Human infant | Male | Berkshire, UK |

## Acknowledgments

This work was supported by funding from: BigC (grant number 18-15R); SDR and LJH – the authors gratefully acknowledge the support of the Biotechnology and Biological Sciences Research Council (BBSRC); this research was funded by the BBSRC Institute Strategic Programme Grant Gut Microbes and Health BB/R012490/1 and its constituent projects BBS/E/F/000PR10353, BBS/E/F/000PR10355, BBS/E/F/000PR10356, and the BBSRC Core Capability Grant BB/CCG1860/1. LJH is supported by a Wellcome Trust Investigator Award 220876/Z/20/Z. For the purpose of Open Access, the authors have applied a CC BY public copyright licence to any Author Accepted Manuscript version arising from this submission.

## Author contributions

Conceptualisation: CAP, LJH, SDR; Formal analyses: CAP, MK, TTK, MR, JM. Investigation: CAP, AN, MK, TTK, NI, WJF, AMM, LM, MR, JAGET, CJB, SAD, JM CJB, RTJ, JAGET; Resources: LJH, SDR; Review and editing: CAP, AN, MK, TTK, NI, WJF, AMM, LM, MR, JAGET, CJB, SAD, JM CJB, RTJ, JAGET, LJH, SDR; Visualisation: CAP, AN, LJH, SDR; Supervision: LJH, SDR; Funding acquisition: LJH, SDR.

## Data availability statement

The raw data supporting the conclusions of this article will be made available by the authors, without undue reservation, to any qualified researcher.

